# Simulating heterogeneous populations using Boolean models

**DOI:** 10.1101/110817

**Authors:** Brian C. Ross, Mayla Boguslav, Holly Weeks, James Costello

## Abstract

Certain biological processes such as cancer development and immune activation are controlled by rare cellular events that are difficult to capture computationally through simulations of individual cells. Here we show that when cellular states are described using a Boolean network model, one can exactly simulate the dynamics of non-interacting, highly heterogeneous populations directly, without having to model the various subpopulations. This strategy captures even the rarest outcomes of the model with no sampling error. Our method can incorporate heterogeneity in both cell state and, by augmenting the model, the underlying rules of the network as well (i.e. mutations). We demonstrate our method by using it to simulate a heterogeneous population of Boolean networks modeling the T-cell receptor, spanning ~ 10^20^ distinct cellular states and mutational profiles.

---

Computer models are widely used to predict behaviors of biological systems, generally by simulating a number of instances of that model and enumerating the range of observed outcomes. Comprehensive simulations of single cells may be realistic given recent progress in constructing elaborate cellular models, but simulation of an entire tissue is far more difficult owing to the vast number of cells, cell types, and their interactions.

As a step towards tissue-scale modeling of cells, we consider the problem of simulating large and heterogeneous populations of *non-interacting* cells. That is, we wish to model the wide variety of cellular states and dynamics experienced by a heterogeneous population of cells in a tissue living in isolation from each other. The output of the approach we propose will be the frequency with which certain events happen over time in a large population. In theory, this result could be obtained by averaging a large number of traditional single-cell simulations spanning the entire population, but in extremely heterogeneous populations it becomes unfeasible to simulate each distinct subpopulation, in which case the traditional recourse is to estimate the population statistics by Monte Carlo (random sampling) [1]. By design, the basic random sampling procedure captures typical outcomes of these simulations, and only rarely finds unusual occurrences. Yet some biological processes are determined by outliers [2], such as rare tumor cells [3–5] or immune clones [6]. We acknowledge that if something is known about the circumstances leading to a rare outcome, one can bias Monte Carlo to oversample that outcome and then correct for the biased sampling (a strategy known as importance sampling [7]), but oversampling inevitably introduces sampling biases.

Here we propose an alternative, exact method for simulating heterogeneous populations, which takes advantage of the observation that discrete models have a finite set of possible states. For these models, one can write the instantaneous state of some individual being modeled using a vector **b**_(*α*)_ of all possible states, having a ‘1’ in the state *α* of the individual and a 0 everywhere else. Assuming deterministic dynamics, the time evolution of this individual can then be written as a linear (matrix) operator *F*_*b*_, and time evolution proceeds by repeated matrix multiplications of the state vector: 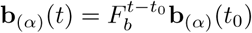 where *t*_0_ and *t* are integer time steps. This approach is always possible in principle for discrete systems, even when it is too computationally expensive to be feasible in practice.

The potential usefulness of a linear representation is that the same equations that simulate an individual *automatically generalize to simulating arbitrary mixed populations* of individuals. The basic idea, as illustrated in Figure 1, is that a population-averaged vector ⟨**b**⟩ = ∑_*α*_*w*_*α*_**b**_(*α*)_ evolves according to the same time-evolution operator *F*_*b*_ as a vector representing an individual, owing to the superposition property of linear systems. We will exploit this fact in deriving a time-evolution operator using an algebra tailored for an individual, and then repurpose that operator to simulate mixed populations.

**Figure 1.**
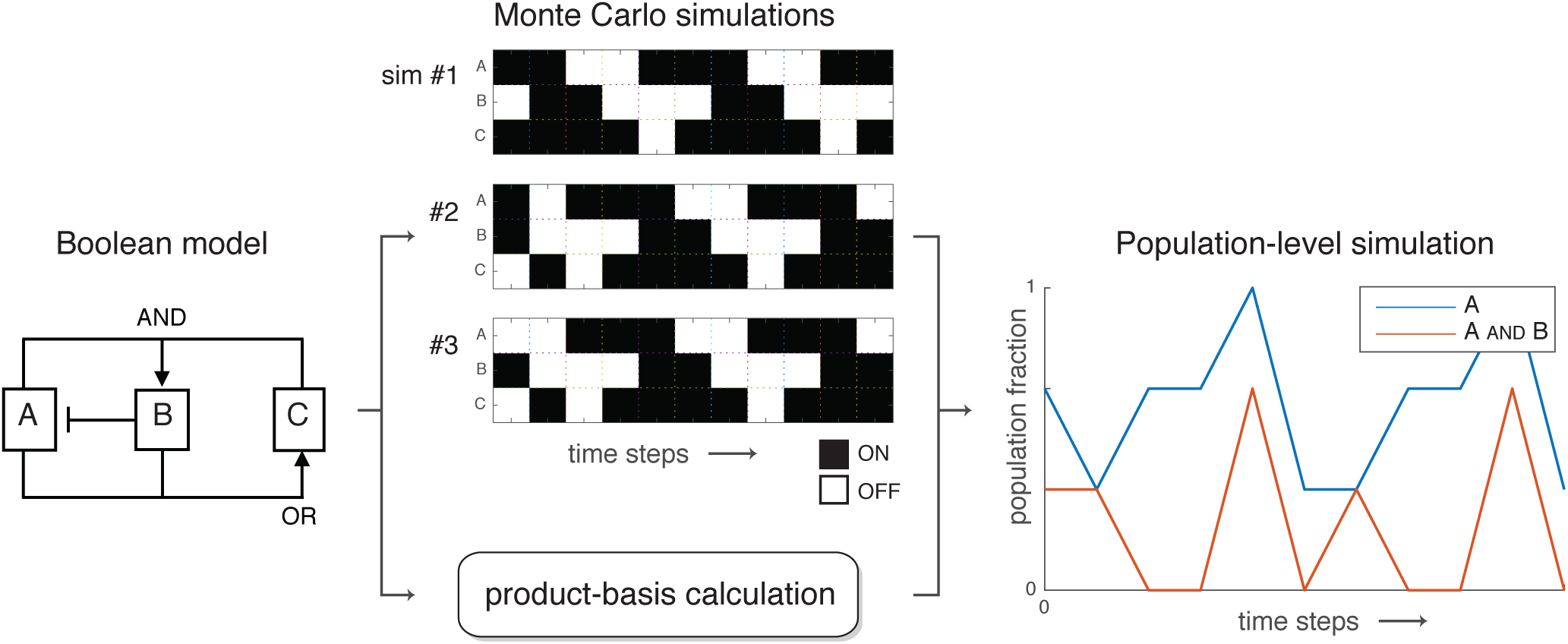
Schematic of a population-level simulation, showing how the population fractions showing activation of variable *A* and variable *A* and *B* evolve over time when the individuals in the population have heterogeneous states. No information about the substructure of the population is lost in the averaging process when one takes into account higher-order correlations (such as *A* and *B*).

The obvious drawback of the linear method of time evolution is the typically huge number of states in a given system, causing both the state vector **b** (or ⟨**b**⟩) and the time evolution operator *F* to be unfeasibly large. For example, Boolean networks are a class of simple models built entirely from on/off variables, yet even these models have an exponential number of states (*2*^*N*^ for *N* Boolean variables in the model). In general a linear representation of the dynamics is typically only feasible if one looks at only a small subspace of the full linear space – i.e. considers only a small subset of the possible set of states of the model. However, this strategy is incompatible with our interest in simulating massively heterogeneous populations: a simulation involving *n*_*s*_ subpopulations necessarily involves at least *n*_*s*_ nonzero entries in both the state vector ⟨**b**⟩ and in the time-evolution matrix *F*. Under this proportional scaling with heterogeneity, a linear representation of the system offers no computational improvement over simple enumerative simulation of each individual subpopulation.

Our proposed solution to the problem of proportional scaling with heterogeneity is to *change linear basis* from the state basis **b** to a new basis **x** in which the size of the *subspace of interest* scales independently of the heterogeneity of the mixed population. (The full spaces of **b** and **x** are necessarily of the same size.) Specifically, the goal of this paper is to introduce a linear basis for Boolean variables that we term a ‘product basis’, and give a prescription for calculating the time-evolution operator in this basis. Although our method only applies to Boolean network models, within the Boolean framework the method is very general, applying to deterministic Boolean models [8], as well as probabilistic [9] and continuous-time [10] Boolean models in the large-population limit. Note that although we assume a synchronous updating rule for all variables in the model, this is without loss of generality because an asynchronous network can be modeled as a probabilistic network with synchronous time steps [11].

By way of comparison, we note the significant advances that have been made in analyzing the possible long-term outcomes, or *attractors* (steady states and limit cycles), of Boolean models [8, 12–19]. Attractors have been found using network-reduction algorithms that find simple networks encoding the long-term behavior of more complex networks [12, 14, 20], methods that solve steady states as zeros of a polynomial equation [21], SAT methods [16, 17, 22], and binary decision diagrams [18, 19, 23] (the introduction of Ref. [13] gives a good review of these techniques). These techniques differ from our proposed method in that they focus on possible long-term behaviors, whereas ours gives an explicit population-averaged simulation of a defined starting population. While our method does have the ability to generate long-term dynamical equations that can be used to find attractors (see Appendix 2), our method differs in that it finds the attractors of a set of characteristics of interest, not the attractors of the complete state.

We first derive the procedure for calculating a product-basis simulation to track transient and long term model behaviors, and give an example calculation illustrating the mathematics. Next, we demonstrate an application of our method by using it to simulate a large heterogeneous population using a published T-cell network [24].

## Methods

Consider a Boolean network model consisting of *N* variables, which updates its state at each time step using a deterministic update rule. We denote the Boolean model variables using Roman letters, e.g. *i*, *j*, …, and use Greek letters to refer to sets of Boolean model variables. For each possible subset of Boolean variables *α* = {*i*, *j*, …}, there is a unique state-basis variable *b*_*α*_ (which we sometimes write as *b*_*ij*…_) and a unique product-basis variable *x*_α_ = *x*_*ij*…_. We consider the state-basis variables to be the components of a vector **b**, and the product-basis variables to be components of vector **x**. There are *2*^*N*^ subsets of all *N* Boolean variables (including both *θ* and the empty subset denoted 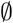), so both **b** and **x** are *2*^*N*^-length vectors.

### Definition 1.

*Consider a Boolean model composed of some set θ of Boolean variables, where B*_*i*_ *represents the state of Boolean variable i* ∈ *θ and follows the convention B*_*i*_ = 1 *for an* on *variable and B*_*i*_ = 0 *for an* off *variable. If κ is the set of* on *variables for some individual, then the values of the state-space and product-space variables describing that individual, indexed by Boolean subset α, are defined by*:

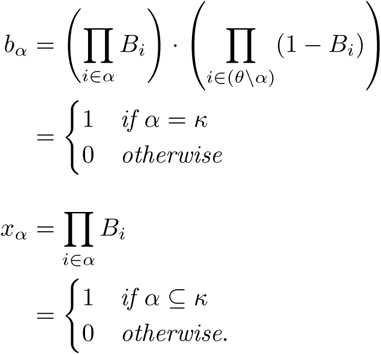

By common convention the empty product 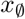 equals 1. The fact that each *x*_*α*_ is a simple product of the values of its subscripted Boolean variables motivates the terminology ‘product basis’. As shown in Appendix 1, a product-basis vector **x** is a fully equivalent representation of a mixed population to the state-basis vector **b**.

### Simulations of mixed populations

We build our simulations by selecting a set of product-basis variables of interest (generally a subset of the full state vector **x**), and associating with each variable *x*_*α*_ an update rule *ƒ*_*α*_ so that *x*_*α*_(*t* + 1) = *ƒ*_*α*_(**x**(*t*)). (The exception is the case of continuous-time Boolean networks in which *ƒ*_*α*_ = *dx*_*α*_*/dt*, but we will treat those separately later.) We construct the simulation in two steps. The first step is to build the single-index update rules *ƒ*_*i*_ (i.e. *α* = {*i*}) for individual Boolean variable *i*, by enumeration of its input states. The second step is to build necessary higher-order update rules *ƒ*_*ij*…_ as needed until the system of equations closes (i.e. until we have solved for each *ƒ*_*α*_ corresponding to a variable *x*_*α*_ appearing in the right-hand side of some other *ƒ*_*β*_). As a necessary preliminary, we first show how to build the change-of-basis operator *T* that converts **b**-basis vectors to **x**-basis vectors.

#### Algorithm 1

Constructing a change-of-basis matrix). *Assume some set of n Boolean variables θ for which κ and α denote subsets. We can construct the change-of-basis matrix T*_*n*_ *that converts a 2*^*n*^ -*length* **b**-*vector indexed by κ to a 2*^*n*^-*length* **x**-*vector indexed by α by assigning a 1 to each element T*_*n*(*ακ*)_ *for which α* ⊆ *κ, and 0 to all other elements*.

*Proof*. Consider the state vector **b**_(*κ*)_ whose entries are all 0 except for a 1 in position *κ*, which describes an individual in state *κ*. The product-space representation **x** of this individual is found by multiplying *x*_*α*_ = ∑_*γ*_*T*_*αγ*_*b*_*γ*_ = *T*_*ακ*_: thus **x** is simply column *κ* of the change-of-basis matrix. Using Definition 1, the value of product-basis variable *x*_*α*_, corresponding to *T*_*ακ*_, is 1 if *α* ⊆ *κ* and 0 otherwise.

We can now give a procedure for calculating the single-variable update rules *ƒ*_*i*_ in terms of its *N*^[*i*]^ Boolean inputs *θ*^[*i*]^. To do so we will work in a reduced space where **b**^[*i*]^ = {*b*_*ρ*_|*ρ* ⊆ *θ*^[*i*]^} and **x**^[*i*]^ = {*x*_*ρ*_|*ρ* ⊆ *θ*^[*i*]^}, as *N*^[*i*]^ is generally small enough that we can explicitly write the change-of-basis operator *T*_*N*^[*i*]^_ in this space using Algorithm 1.

#### Algorithm 2

(Computing *ƒ*_*i*_). *Define* **k**^[*i*]^ *as the vector such that* 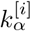 *is 1 if Boolean input pattern* 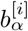 *produces a 1 in output Boolean variable i, and 0 otherwise. Then* 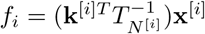, *which is a linear equation in* **x**^[*i*]^ ⊆ **x**.

*Proof*. By definition *ƒ*_*i*_ = **k**^[*i*]*T*^**b**^[*i*]^, as this expression reproduces the output rule. Using the fact that *T*_*N*^[*i*]^_ is invertible (proved in Appendix 1), we write 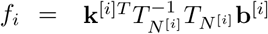 and identify *T*_*N*^[*i*]^_**b**^[*i*]^ = **x**^[*i*]^, which proves the formula.

Using the set of single-index *ƒ*_*i*_, one can compute the linear time-evolution function of any multi-index product-basis variable of interest *ƒ*_*ij*…_ using the following method.

#### Algorithm 3

(Computing *ƒ*_*ij*…_). *We first compute ƒ*_*α*_ *for α* = {*i*, *j*, …} *as ƒ*_*α*_ ← *ƒ*_*i*_(**x**) · *ƒ*_*j*_(**x**) · … *(expressed in terms of* **x**-*basis variables). Next we distribute each term inside the product, so that ƒ*_*α*_ *is a weighted sum of products of* **x**-*basis variables, and replace each nonlinear product of terms x*_*β*_ · *x*_*γ*_ · … *appearing inside ƒ*_*α*_ *with the product-basis variable x*_*μ*_ *where μ* = *β* ⋃ *γ* ⋃ …. *This gives an expression for ƒ*_*α*_ *that is linear in* **x**.

*Proof*. First we show that *f*_*α*_ = *ƒ*_*ij*…_ equals the product *ƒ*_*i*_ · *ƒ*_*j*_ · ….

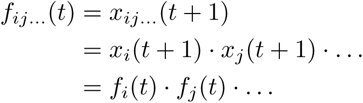

Since this always holds true irrespective of the time step *t*, we simply calculate *ƒ*_*ij*…_ = *ƒ*_*i*_ · *ƒ*_*j*_ · ….

The second step is to show that each *x*_*β*_ · *x*_*γ*_ · … equals *x*_*β*⋃*γ*⋃…_. Let *k*, *l*, … be the elements of *μ* = *β* ⋃ *γ* ⋃ …. Then 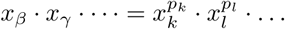 where *p*_*k*_, *p*_*l*_ ··· ≥ 1 are the respective number of times *k*, *l*, … appear in *β*, *γ*, …. Since all *x*_*i*_ are Boolean variables, we have 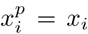 for any *p* ≥ 1. Thus *x*_*β*_ · *x*_*γ*_ ···· = *x*_*k*_ · *x*_*l*_ ···· = *x*_*μ*_.

Using Algorithms 2 and 3, we can give the full procedure for building a simulation that time-evolves any arbitrary set of product-basis variables of interest describing some individual modeled by the Boolean rules. We denote the set of variables of interest as Ω_0_. Our algorithm constructs *ƒ*_*α*_ for each *α* ∈ Ω_0_, then additional *ƒ*_*β*_ to time-evolve each *x*_*β*_ appearing in the right-hand side of the *ƒ*_*α*_, etc. until the equations form a closed system.

#### Algorithm 4

(Building a product-basis simulation). *First we compute the full set of single-index ƒ*_*i*_ *using Algorithm 2. Next we initialize the total set of required product-basis variables* Ω ← Ω_0_, *and the set of product-basis variables whose dynamics we have solved* 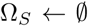. *Finally we iteratively solve ƒ*_*α*_ *for each α* ∈ Ω\Ω_*S*_ *using Algorithm 3, while updating* Ω_*S*_ ← Ω_*S*_ ⋃ {*α*} *and* Ω ← Ω ⋃ {*μ*} *for each variable x*_*μ*_ *appearing on the right-hand side of ƒ*_*α*_. *Iteration continues until* Ω_*S*_ = Ω.

*Proof*. We first note that Ω eventually converges to a finite set owing to the fact that the full variable space *θ* is a finite set, and that Ω ⊆ *θ* only gains, never loses, elements of *θ* at each iterative step. Since a) Ω_*S*_ ⊆ Ω, b) Ω_*S*_ accumulates 1 term in Ω\Ω_*S*_ at each iterative step, and c) Ω converges, the algorithm always ends in a finite number of steps. When the algorithm ends the set of solved equations Ω_*S*_ contains the set of variables appearing in the right-hand side of those equations Ω_*R*_ ⊆ Ω = Ω_*S*_, so the resulting system of equations is closed.

Writing our final closed system of linear equations as a square matrix *F*, we have a very simple update rule for simulations using the product-basis variables: **x**(*t* + 1) = *F* · **x**(*t*).

Our final step is to generalize to a mixed-population simulation.

#### Definition 2.

*Define a population-level state vector* ⟨**x**⟩ *as a population-weighted linear combination of the state vectors of the subpopulations* **x**_(*α*)_:

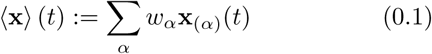

*where the weighting factors w*_*α*_ *are proportional to the occurrence of subpopulations α in the overall mixed population*.

Note that we are free to choose the normalization constant *n* = ∑_*α*_*w*_*α*_. In all our examples we will choose *n* = 1, giving *w*_*α*_ the interpretation as the population fraction of state *α*. Irrespective of the choice of *n*, this representation can be used to time-evolve mixed populations using our existing operators.

#### Claim 1.

*Each element* ⟨*x*_*α*_⟩ *of vector* ⟨**x**⟩ *is proportional to the overall occurrence of characteristic α in the overall mixed population. For n* = 1, ⟨*x*_*α*_⟩ *is the average occurrence of that trait in the mixed population*.

*Proof*. Equation 0.1 is the definition of a **w**-weighted average of characteristics **x**, and **w** is proportional to the population fraction.

#### Claim 2.

*Any linear time-evolution operator that properly time-evolves an individual in any arbitrary state* **x**_*α*_ *also correctly time-evolves any arbitrary mixed population of individuals* ⟨**x**⟩.

*Proof*. This follows from the fact that *F* commutes with the sum over subpopulations.

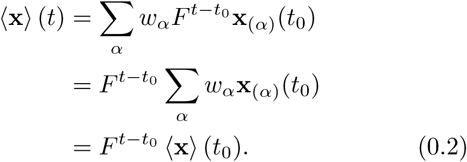

Thus the time-evolution operator produced by Algorithm 4 correctly time-evolves the mean occurrence of the characteristics ⟨**x**⟩ in any mixed population, *despite the fact that this operator was derived for an* **x** *describing an individual* (notably in assuming that each *x*_*α*_ is Boolean).

#### Example 1: building equations

Suppose we wish to follow the time evolution of variable *A* in the model shown in Figure 2. The first step is to compute the change-of-basis matrices that will be used to compute the single-variable update rules *ƒ*_*A*_, *ƒ*_*B*_ and *ƒ*_*C*_. Variable *A* takes input from the single variable *B*, so calculating *ƒ*_*A*_ requires the change-of-basis matrix *T*_1_. Ordering the elements by the subscripts 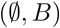 respectively, and applying Algorithm 1, we obtain

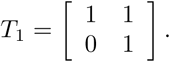

Variables *B* and *C* each take input from two variables. Ordering *B*’s input states 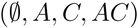, and C’s input states 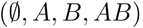, we find that both *ƒ*_*B*_ and *ƒ*_*C*_ are computed using

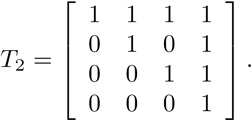

Next we build the single-index update functions. Variable *A* takes input only from variable *B*, so the possible patterns of active inputs are 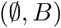, corresponding to the state-basis variables 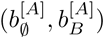. The respective outputs are (1, 0) = **k**^[*A*]*T*^ due to the not gate, from which we can immediately calculate *ƒ*_*A*_ using Algorithm 2.

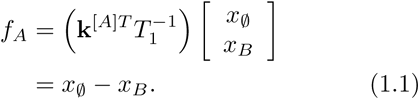

In the same way we find that the input patterns 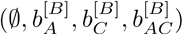 to variable *B* lead to outputs **k**^[*B*]*T*^ = (0, 0, 0, 1), and the input patterns 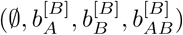 to variable *C* lead to outputs **k**^[*C*]*T*^ = (0, 1, 1, 1). From these and the 2-variable change-of-basis matrix *T*_2_ we calculate

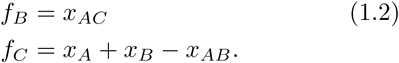

Having built the single-index update functions, we can now derive a linear system that tracks the time evolution of variable *A*. The immediate equation for this purpose is *ƒ*_*A*_ which we already derived (Equation 1.1), but since it involves *x*_*B*_ and 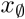 our simulation must also track those variables through time using Equation 1.2 along with

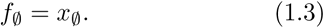

Equation 1.2 requires that we track a new multi-index variable *x*_*AC*_, requiring us to solve *ƒ*_*AC*_ using Algorithm 3.

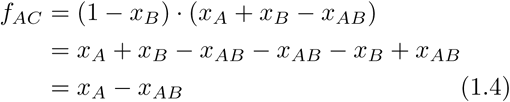

We continue the process of identifying new variables and solving for their update functions until the equations form a closed system.

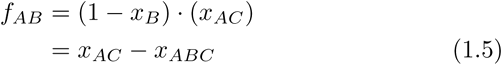

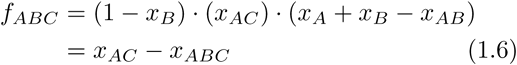

Equations 1.1-1.6 together with an initial state in (*x*_*A*_, *x*_*B*_, *x*_*AC*_, *x*_*AB*_, *x*_*ABC*_) describe the time evolution of these quantities in a single Boolean network as a sequence of 0s and 1s in each variable. The final step is to reinterpret these equations as describing the dynamics of a mixed population of networks, formally by taking the mean of both sides of each equation.

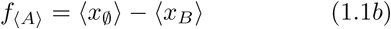

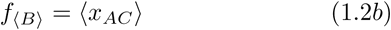

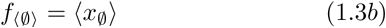

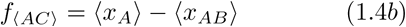

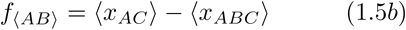

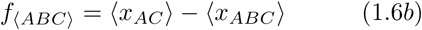

The angle-brackets denote an average, and we have used the notation ⟨*x*_*i*_(*t* + 1)⟩ = *ƒ*_⟨*i*⟩_ Per Claim 2 the linear equations are unaffected by the averaging process, so the same equations used to derive the dynamics of a single instance of a model also describe the mean values of those same variables in a mixed population. Whereas the product variables take on binary values for an individual, the population-averaged variables are real-valued on the interval [0, 1] (using our normalization): for example we would set ⟨*x*_*A*_⟩ = 0.4 if 40% of the population has gene *A* set on.

**Figure 2.**
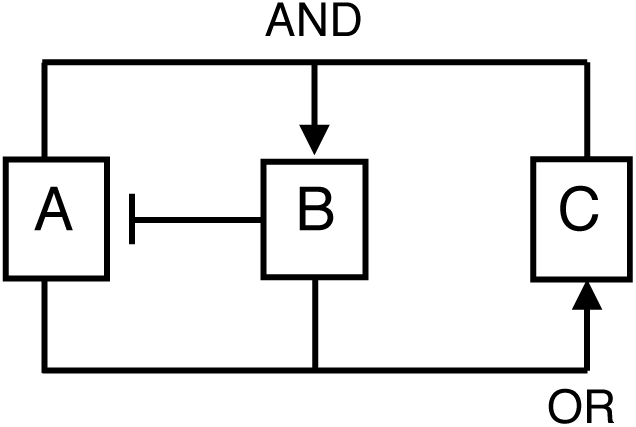
The 3-Boolean network used in Example 1. Arrows indicate the update rules for each variable: for example if either A or B is on at time *t* then C will be on at time *t* + 1; otherwise C will be off.

### Probabilistic and asynchronous Boolean networks

Our method can be applied to probabilistic Boolean networks (PBNs) [9, 25], in which several state transitions are possible at each time step with different probabilities. As we will show, our algorithm works as given in the *large-population limit* in which time evolution of the average state ⟨**x**⟩ becomes deterministic, despite the fact that each individual PBN is stochastic.

Applying our method to PBNs requires that we reinterpret the meaning of the time-evolution equations. For an individual we write:

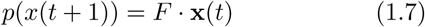

where *p*(·) denotes the likelihood of an outcome. With this new interpretation of the time-evolution operator *F*, our method still works, although we note here several changes to the logic. 1) In Algorithm 2 we generalize each **k**^[*i*]^ to be a vector of likelihoods that each respective input pattern produces a 1 in the output variable, so that *ƒ*_*i*_ = **k**^[*i*]*T*^**b**^[*i*]^ as before. 2) The multiplication rule, now reading *p*_*x*_*ij*…__ = *p*(*x*_*i*_)*p*(*x*_*j*_) …, used by Algorithm 3 still holds, owing to the independence of outcomes *p*(*x*_*i*_), *p*(*x*_*j*_), etc. 3) We point out that although *p*(**x**) is real-valued, the state vector of an individual **x** is still binary, so 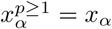 as before (Algorithm 3).

Despite the fact that our algorithm produces valid product-basis equations of the form 1.7, the resulting linear system of equations does not form a closed system, simply because the left-hand side uses different variables than the right (*p*(**x**) versus **x**). Our method therefore cannot be used to simulate an individual instance of a PBN. However, by averaging both sides and taking the large-population limit so that *p*(**x**) → ⟨**x**⟩, the system closes and reproduces Equation 0.2. Thus our system of equations accurately tracks the deterministic dynamics of arbitrary mixed populations of PBNs in the infinite-population limit, despite being unable to model stochastic individuals.

Large populations of asynchronous networks behave identically to large populations of PBNs [11] if we define a uniform time step: the likelihoods of the various possible updates over that time step give the state-transition weights in the corresponding synchronous PBN. If this time step is small enough, then the likelihood of two causally-connected asynchronous updates happening in the same step is small, and in this limit the local update rules for a PBN accurately model the asynchronous network. Therefore our analysis also applies to large populations of asynchronous networks for small time steps.

### Continuous-time Boolean networks

Probabilistic networks give one way to incorporate rate information into our model; another way is to work in continuous time using differential equations [10]: *ƒ*_*A*_ = *dx*_*A*_/*dt*. The differential form does require one change in our method: the rate of change of a higher-order variable is found by using the product rule of derivatives. Whereas under a discrete update *ƒ*_*ABC…*_ is the product *ƒ*_*A*_ · *ƒ*_*B*_ · *ƒ*_*C*_ · …, for the differential case we compute:

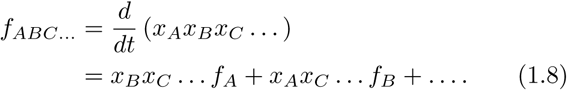

Also, under discrete updates the trivial function is 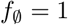 but with differential updates it is 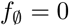.

## Results

### Product-basis equations recapitulate Monte Carlo simulations

We tested the product-basis method using 10^4^ randomly generated 10-node deterministic networks, where each node’s input combined the output of 1-4 randomly-selected nodes using randomly-generated logic rules. For each network, we ran 100 Monte Carlo individual simulations using a random ensemble of initial states (equal probability for all states), and compared the population average of certain state variables with a product-basis simulation using the same starting population distribution. In each case, the product-basis simulation reproduced the average of the Monte Carlo simulations. Next, we ran 10^4^ tests of probabilistic networks (PBNs), again using 100 Monte Carlo runs per test. In order to generate realistic PBNs, we augmented the original time-evolution functions 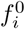 with random rate parameters *r*_*i*_, leading to the equations 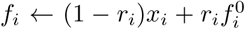. Again the equations of the product-basis method reproduced the results of Monte Carlo simulations, this time to within sampling error 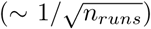. Finally, we tested 2.5 × 10^3^ continuous-time networks generated using the same rule (which are slower to check by Monte Carlo), and again found complete agreement with Monte Carlo.

### Product-basis method can simulate highly heterogeneous populations beyond the scope of Monte Carlo

To demonstrate the product-basis method on very heterogeneous populations, we applied it to the T-cell activation network described in Ref. [24] (see Figure 10 and Table 2 of that paper). The T-cell network is a deterministic, 40-node network with fifty-four edges, multiple feedback loops, and whose attractors include both steady states and limit cycles. For demonstration purposes, we show a traditional Monte Carlo simulation of an *individual* T-cell network in Figure 3A, obtained by choosing an initial Boolean state and straightforwardly applying the model rules over each successive time step. This particular simulation shows transient (nonrepeating) behavior for the first 10 time steps, leading to a limit cycle with a repeating period of 6 time steps.

**Figure 3.**
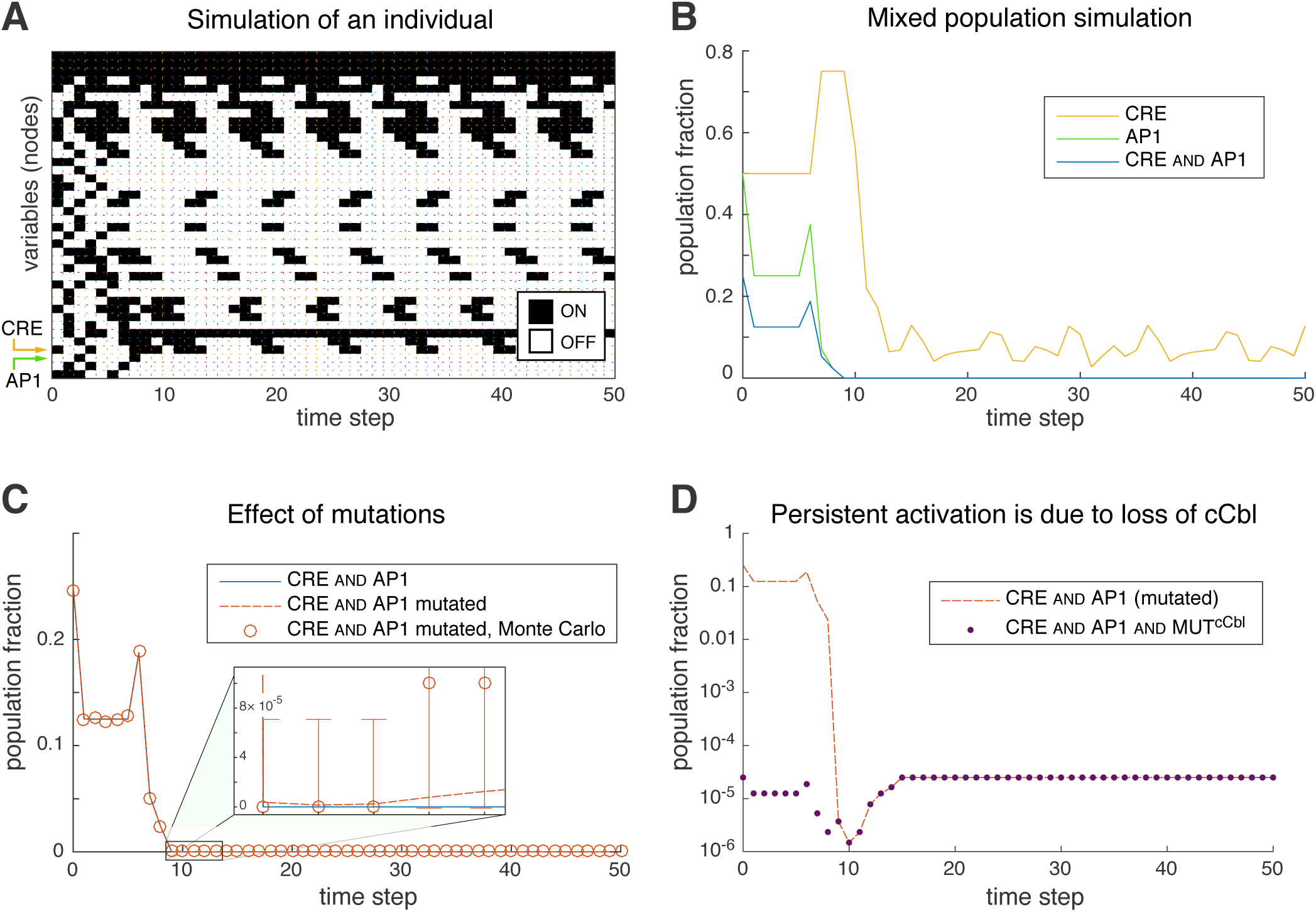
**A)** Time evolution of one instance of the T-cell network [24], starting from a random initial state. White/black rectangles signify off/on Boolean states. **B)** Time evolution of the population fraction having activated CRE elements and/or expressing the transcription factor AP1 in a heterogeneous population of T-cell networks, computed using a product-basis calculation. The heterogeneous population begins at *t* = 0 as a uniform mixture of all possible 2^33^ ≈ 10^10^ initial states of the upstream portion of the model. **C)** The effect of a 10^−4^ knock-out mutation rate per gene in the heterogeneous population. Monte Carlo, but not the product-basis calculation, required this high rate of mutations in order to detect persistent coactivation of CRE and AP1. **D)** The co-occurrence of CRE activation and AP1 expression in mutated networks shown on a log-10 scale (dotted red line), compared with the amount of this coexpression coincident with mutated cCbl (purple dots). The mutated fraction was computed by subtracting the time series of CRE and AP1 and WT-cCbl from the time series of CRE and AP1.

Next, we performed a population-level simulation using the product-basis method. We first provided a target set of variables to follow in time, which the product-basis algorithm used to generate a set of time-evolution equations involving those variables (along with other variables that were added automatically to close the system of equations). We chose to track three variables: cyclic-AMP response element (CRE) mediated gene activation, the AP1 (Activating protein 1) transcription factor, and their co-occurrence which we label CRE and AP1. The cooccurrence variable was not redundant: although CRE and AP1 is simply the product CRE × AP1 for an individual (because the variables are Boolean), this is *not* true at the population level. (For example if the levels of CRE and AP1 are both 0.5, then the level of CRE and AP1 could equal 0.25 if CRE and AP1 states are uncorrelated in the population, 0 if they are perfectly anticorrelated, 0.5 if they are perfectly correlated, or any other value in between.)

We first generated the product-basis time-evolution equations for our three variables of interest. Next, we set the model variables to an initial state representing a mixed population, and used the time-evolution equations to track the population-level average of each of the three variables for 50 time steps (Figure 3B). We stress that this is an exact result, with no sampling error. The starting population we considered was a uniform mixture of all possible 2^40^ initial states of the Boolean network, but because 7 of these variables are not upstream of either CRE or AP1, we consider our effective initial population size to be 2^33^ ≈ 10^10^. At this level of diversity, it would require extensive computational resources to reproduce our exact result by exhaustive enumeration over the initial states.

Next, we demonstrated the ability of the product-basis method to analyze mutations in the network by including the full set of possible gene knock-outs in the T-cell activation network. We did this by adding a set of ‘wild-type’ variables to the network, one for each original variable in the system, and included the wild-type variables in the update rules using an and operation. For example, an update rule reading [*A* ← *B* or *C*] became [*A* ← (*B* or *C*) and *A*^WT^]. Note that the presence of mutations effectively doubles the size of the network and thus vastly increases the heterogeneity, which is determined by both the number of activation states of the original variables and the number of mutational profiles, in total spanning the order of 10^20^ different subpopulations. Enumeration over the initial states is impossible at this level of diversity, and the traditional solution is a sparse sampling method such as Monte Carlo, although that loses the ability to resolve very rare subpopulations. Despite this heterogeneity, our product-basis method was able to produce the exact time-evolution equations for CRE and AP1. We chose an initial population that contained each possible combination of knock-out mutations at a 0.01% mutation rate per variable (roughly the highest possible rate of homozygous knockouts given a 1%-per-gene-locus mutation rate [26]) superimposed upon a uniform mixture of each activation state (adjusted so that mutated genes always began off). From this initial state, we again followed the exact time course of the CRE and AP1 population fraction, and compared it to our original wild-type result (Figure 3C). Notably, a small fraction of the population reached a steady state showing both CRE activation and AP1 expression. We also validated the result (to within statistical error) using Monte Carlo, although as shown in the figure, Monte Carlo was only useful for comparing the transient behavior, not the rare persistent subpopulations which fell far below its sampling resolution.

Finally, we examined the mutations leading to CRE and AP1 coexpression. We hypothesized that this was due to knockout of the cCbl gene in the recurrent core of this network, and tested this hypothesis by generating the time course of the three-way co-occurrence of CRE and AP1 and WT-cCbl, where WT denotes the respective wild-type variable. This final time series dropped to zero at steady state, indicating that mutations in this variable are necessary for persistent CRE and AP1 coexpression (see Fig. 3D), and that no other set of knockout mutations could recapitulate this phenotype.

Our results from the T-cell network demonstrate several important aspects of our method. First, we are able to simulate extremely heterogeneous populations, involving far more subpopulations than could be analyzed individually. Second, although our method only deals with heterogeneity in the states of the Boolean variables, we can still simulate a genetically-heterogeneous population by augmenting the Boolean network with mutation variables. Third, we can exactly model subpopulations that are present at very low levels, which are difficult to resolve by random sampling (see the error bars in Figure 3B). For example, the contribution of each triple-mutant was factored in even though a given triple-mutant was present in only 10^−12^ of the population. While one might artificially raise the Monte Carlo mutation rate to oversample the mutations [7], this has the disadvantage of overweighting the effect of multiple mutants, even though realistic evolutionary paths take one or very few mutational steps at a time [27]. In contrast, our exact result is dominated by the evolutionarily-accessible subpopulations that are closest to wild-type.

### MATLAB code

The code used to generate these results is named `tCellActivationEx.m`, and is available for download at https://github.com/CostelloLab/ProductBasis. The equation-generating process for Figures 3B and 3C took ~ 3 and ~ 300 seconds respectively using our code (written in MATLAB R2015b 8.6.0.267246, running on a 2.6GHZ Intel core i7 Mac with OS 10.9.5). The Monte Carlo comparison in Figure 3C (*n*_*runs*_ = 10^4^) took ~ 140 seconds.

## Discussion

Our product-basis method allows the direct simulation of highly heterogeneous populations, including the transient processes that are ignored by Boolean-attractor analyses. Our approach can be used to follow single variables of the system over time, as well as the correlations between these variables that are both necessary and sufficient to fully describe the dynamics of the population. We also showed that our method, when applied to a network augmented by mutation nodes, can effectively explore heterogeneity in the network rules in addition to heterogeneity in network state. In each of our simulations, all subpopulations are exactly accounted for in the output time series, no matter how rare. The only requirement for use of our approach is that the underlying model be built using Boolean variables.

The key to our method is to write the time-evolution equations as a linear system, but in a different basis than the usual state space basis. Our variables have several advantages over state space variables. First, descriptors of a mixed population naturally use words that correspond more closely to our variables than to individual states. For example, we might specify that half the population starts with both genes *A* and *B* on, which implies that *x*_*AB*_ = 0.5, but is agnostic about the state of the rest of the network. Another advantage is that our equations often close using relatively few of our product variables for any mixed population, whereas the number of equations required in the state space basis scales with the heterogeneity of the population: the simulations we showed in Figure 3 would require all 2^40^ state space variables. Thus our choice of variables is better for modeling very heterogeneous populations. Finally, our basis allows for a variable-factorization scheme which can help to simplify the calculation if we only care about the long-term behavior.

We acknowledge that our method can become intractable for complex networks due to the fact that construction of these simulations is potentially an exponential problem. Fully-connected Boolean networks with random logic rules will probably always be challenging, but we believe it should be possible to improve our performance on certain network motifs such as downstream feedback loops that give our method difficulty. Our proposed basis is only one of very many obtained through shear-type transformations similar to our *T* matrices, and it may well be that other choices of basis perform better on certain types of network. Future work will explore this possibility. Additionally, the equation-reduction method for finding attractors (Appendix 2) is somewhat of an art, and our future work will aim to improve this part of the calculation for typical network models (although the attractor analysis is also known to be NP-hard [28]).

The product-basis method can be applied to any system involving heterogeneous populations, as long as the individuals in a population can be modeled using Boolean logic. Heterogeneity plays a major role in such varied systems as healthy and cancerous tissues, evolution at the organism scale, and the social dynamics of unique individuals [29]. In all of these cases, rare and unexpected dynamics are difficult to capture by simulations of individuals, while pure attractor analyses may miss important aspects of the dynamics. We have demonstrated that the methodology outlined here can help to capture these important but elusive events.

## Appendices Appendix 1: product variables form a complete and independent basis spanning the state space

### Claim 3.

*The change-of-basis matrix carrying* **b** *variables to* **x** *variables is an invertible transformation*.

*Proof*. Our proof first requires us to explicitly write the change-of-basis matrix *T*. We can do this using Algorithm 1 once we have chosen a particular ordering of the entries of both the **b** and **x** vectors. Here we will choose to order the entries of both **b** and **x** by binary order of the subscripts on their respective state variables. In other words, the index is given by the binary number having 1s in the positions indicated by the subscripts of each state variable: the state variable written as *b*_*α*_ = *b*_*ij*…_ is the *m*th entry in the state-space vector **b** and *x*_*β*_ = *x*_*ij*…_ is the *m*th entry in the product-space vector **x**, where *m* = 2^*u*-1^ + 2^*v*-1^ + · · · + 1 if we take 1 ≤ {*i*, *j*, …} ≤ *N*. This choice of ordering fixes a particular representation of the change-of-basis matrix. For example, for the case of three Boolean variables the change-of-basis matrix is:

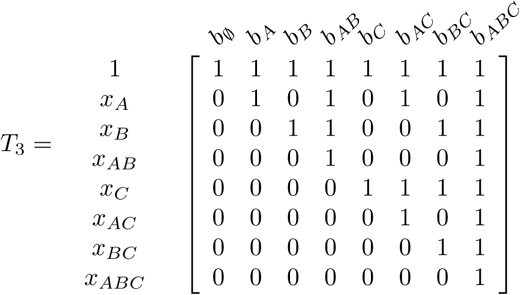

Our objective is to show that for arbitrarily-sized systems, the change-of-basis matrix is upper-triangular, with ones on the diagonal.

#### Lemma 1.

*The diagonal entry x*_(*m*)_ *equals 1*.

*Proof*. The vector component *x*_(*m*)_ lying along the diagonal *T*_*mm*_ is the value of product variable having the same set of indices as *b*_*ij*…_, namely *x*_*ij*…_. Since *i*, *j*, … are the set of on variables, *x*_*ij*…_ evaluates to *B*_*i*_ · *B*_*j*_ · · · · = 1 · 1 · · · · = 1.

#### Lemma 2.

*All entries below the diagonal (x*_(*p*)_ *for p* > *m) equal 0*.

*Proof*. Assume we have some *x*_(*p*)_ ≠ 0 for *p* ≠ *m*, corresponding to some product-space variable *x*_*kl*…_ which is a product of Boolean variables *B*_*k*_ · *B*_*l*_ · … which must each be nonzero in order for *x*_(*p*)_ = *x*_*kl*…_ to be nonzero. Since {*B*_*i*_, *B*_*j*_, …} is the set of oN Boolean variables, we have that *P* = {*k*, *l*, …} is a proper subset of *M* = {*i*, *j*, …}; note that *P* ≠ *M* because *p* ≠ *m* and each index position corresponds to a unique set of Boolean variables (because binary representations of numbers are unique). Thus 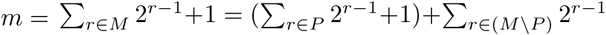 where *M\P* is nonempty, implying that *m* > *p*. Therefore, if *p* > *m* then the premise cannot hold, and *x*_(*p*)_ = 0.

Repeating this logic over all indices *m*, we conclude that each diagonal entry of the change-of-basis matrix is 1, and the entire lower triangle is 0. Since the change-of-basis matrix is triangular the eigenvalues are the diagonal entries, which are all 1. Therefore the determinant is 1, so the change-of-basis transformation is non-degenerate (and volume-preserving) and therefore invertible.

## Appendix 2: equation-reduction for longtime behavior

A mixed-population simulation may or may not be practical, depending on whether the system of linear equations closes with a manageable number of variables. One way to simplify the equations is to focus on the long-time behavior of the system, after quickly-decaying eigenmodes have dropped near zero and time evolution effectively occurs in the smaller subspace of slowly-decaying or persistent eigenmodes. In the limit where we focus on the infinite-time behavior, this subspace is simply the attractors of the system (steady states or limit cycles). Our general strategy will be to 1) identify fast-decaying modes as they appear in *F*, set them to zero and use the resulting expression as a constraint that can be used 2) eliminate variables from later steps of the calculation. In order to be useful for simplifying calculations, these steps must be taken before the linear system closes – in other words, when our partially-computed update function *F*_*partial*_ is still a rectangular matrix.

Our strategy for finding decaying modes in *F*_*partial*_ relies on a QR decomposition, which can be performed on rectangular matrices. In the case of a deterministic network, all eigenvalues have modulus 0 or 1 implying that all decaying modes are strict linear dependencies. (*Proof*: The state space and thus by Appendix 1 the product-basis space is finite, so every eigenmode must either decay to zero in finite time implying that λ = 0, or else repeat in Δ*t* time steps so that |λ| = 1 and arg(λ) = 2*πk*/Δ*t* for integer *k*.) The set of linearly-dependent rows in *F* is found by identifying (numerically) zero entries along the diagonal of the *R* matrix. The dependencies themselves are then calculated by solving the matrix equation *N* = *CD* for polynomial coefficients *C*, where *N* and *D* are the submatrices of *F* containing its independent/dependent rows. For PBNs, the complication is that decaying eigenvalues of the full system are generally nonzero, so the decaying modes are not simple linear dependencies. Note that each decaying eigenvalue is part of the spectrum of *F*_□_ (the square component of *F*_*partial*_), although the reverse is not true. Our implementation uses the spectrum of *F*_□_ as a set of candidate eigenvalues, and for each sufficiently-small candidate λ (based on a user-defined threshold) finds the linearly dependent subspace using a QR decomposition of *F*_*partial*_ - λ*I*_*partial*_, where *I* is rectangular with ones on the diagonal and zeros elsewhere.

Each constraint takes the form of a polynomial in **f** that effectively becomes zero after a certain number of time steps have passed, implying that the same polynomial in **x** becomes zero one time step later. The general form of a constraint is thus:

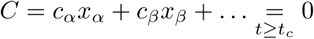

There are several ways in which these constraint polynomials could help simplify the calculation, but we must be careful that 1) application of a constraint should not undo the work of a prior constraint, and 2) there should be a way of resolving how two constraints attempt to remove the same variable. Here we give two constraint algorithms that help to simplify long-time analyses in the cases of probabilistic and deterministic Boolean models respectively.

### Algorithm 5

(Equation-reduction method 1). *Given constraint C, we choose the ‘lowest’ variable x*_*ϕ*_ *(which may be* 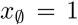) *based on a binary representation of its indices phi as in Appendix 1, and eliminate that variable from all current and future time-evolution equations in F by simple substitution: x*_*ϕ*_ → -(*c*_*α*_/*c*_*ϕ*_)*x*_*α*_ -(*c*_*β*_/*c*_*ϕ*_)*x*_*β*_ - …. *Constraining a time-evolution equation valid from time t_e_ using this formula yields a different time-evolution equation valid for t* ≥ max(*t*_*e*_, *t*_*c*_).

*Proof*. The substitution is justified for *t* ≥ *tc*, so the resulting time-evolution equation is accurate after time *t*_*c*_. A given constraint either eliminates a variable (if the constraint polynomial contains only one term), or else replaces a variable exclusively with ‘higher’-ordered variables. Each replacement with higher-ordered variables increases the total number of indices in each constrained term by at least 1, and since the maximum number of indices is *N* the process of applying constraints eventually ends. Each constraint application permanently eliminates one variable from the finite 2^*N*^-sized variable space, so the process of constraining the equations eventually ends, which is only possible if there are no more fast-decaying modes. Thus this process eventually eliminates the fast-decaying eigenspace, while preserving the slowly-decaying/persistent eigenspace (because the constrained *ƒ* equations are valid after a certain time *t*_*c*_*max*__).

We use reduction method 1 for PBNs and asynchronous networks as the second method does not apply to them. This method is relatively inefficient since it only eliminates one variable per constraint. There is also some indeterminism: if two constraints apply to a given term in an equation, we simply choose one of the two constraints to apply. We give a simple demonstration of reduction-method 1 (on a deterministic network) by using it to find the attractors of Example 1.

### Example 1, continued

Eqs. 1.1-1.6 in Example 1 contain a single linear dependency: *ƒ*_*AB*_ = *ƒ*_*ABC*_. Since the network is deterministic, this dependency guarantees that after the first time step *x*_*AB*_ will equal *x*_*ABC*_, so we write 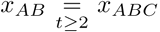. We use this fact to eliminate *x*_*AB*_, giving a new set of steady state equations:

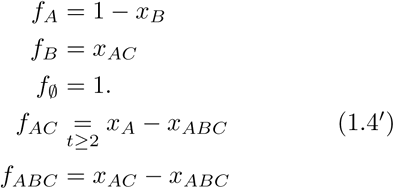

Our new set of equations has the same non-zero eigenspace as the original set (1.1-1.6), except Eq. 1.4′ is only valid from the second time step onwards. However, the new equations lack the null eigenspace because we removed the only linear dependency. The 5 eigenvalues all have phases that are multiples of 2*π*/5, indicating that the sole attractor is a limit cycle with a period of 5 time steps. We acknowledge that in this case the constraint helps find the attractors but does not simplify the calculation, since it only appears after the full *F* has been calculated.

### Algorithm 6

(Equation-reduction method 2). *Given the constraint polynomial C, we select a variable x*_*ϕ*_ *whose indices ϕ have the property* 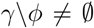 *for every other term x*_*γ*_. *Constraining a time-evolution equation of a deterministic model valid from time t*_*e*_ *involves constraining every variable x*_*ρ*_ *in that equation that is factorized by x*_*ϕ*_, *in the sense that* 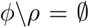 *Application of the constraint takes x*_*ρ*_ → -(*c*_*α*_/*c*_*ϕ*_)*x*_*α*⋃*ρ*_ - (*c*_*β*_/*c*_*ϕ*_)*x*_*β*⋃*ρ*_ - …, *yielding an equation that is valid for t* ≥ max(*t*_*e*_, *t*_*c*_).

*Proof*. For time *t* ≥ max(*t*_*e*_, *t*_*c*_) both the original time-evolution equation and the constraint equation are valid, implying that *x*_*ϕ*_ = -(*c*_*α*_/*c*_*ϕ*_)*x*_*α*_ - (*c*_*β*_/*c*_*ϕ*_)*x*_*β*_ - …. For deterministic models we have *x*_*ρ*_ = *x*_*ϕ*_ · *x*_*ρ*_, since *ϕ* ⊆ *ρ* and indices can be freely duplicated in deterministic models. Substituting the two formulas gives the constraint rule. Since each constraint application either eliminates *x*_*ϕ*_ or replaces it with variables containing more indices, the arguments of constraint method 1 apply, showing that the method eventually eliminates the fast-decaying eigenspace while leaving correct time-evolution equations for the slower-decaying modes.

Equation-reduction method 2 is considerably more aggressive than method 1, because each constraint eliminates not only one or more variables involved in the constraint, but also any variables that are factorized by them: thus a dependency involving a low-order variable with few indices can exclude a significant fraction of the variable space. We demonstrate this method 2 using a second example.

### Example 2: applying constraints

Suppose we want to find the long-time behavior of Boolean variables A, C, and E in the network of Figure 4. After two iterations of solving for time-evolution functions we have:

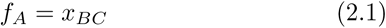

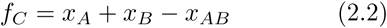

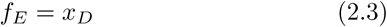

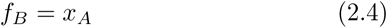

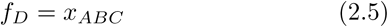

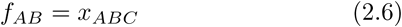

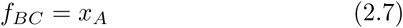

At this point there are two linear dependencies: *ƒ*_*AB*_ = *ƒ*_*D*_ and *ƒ*_*BC*_ = *ƒ*_*B*_, implying that

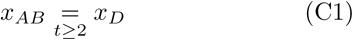

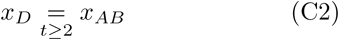

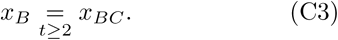

Since we are only interested in the long-time behavior, we will use these constraints to simplify our equations. For example, if we were to retain *x*_*ABC*_ it would be affected by the relationships involving *x*_*AB*_, *x*_*B*_, and *x*_*BC*_, and it would not be possible to enforce all of these by substitution because there is only one *B*-index on *x*_*ABC*_. But we can simultaneously enforce all constraints by *multiplying* them:

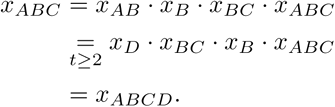

More generally, the first constraint attaches an *AB* index to every variable containing a *D*, and a *D* index to each variable with *AB* indices, and the second constraint adds a *C* index to every variable with a *B* index.

Constraining our system and eliminating disused variables gives us

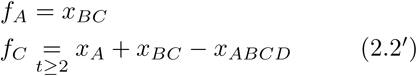

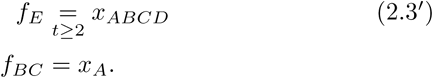

Our new equations require us to solve for another variable (while applying the constraints):

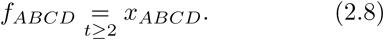

The system is now closed (5 equations involving 5 variables), so if our goal is to produce a simulation (valid from time step 2 onwards) then we are done. However, if our objective is to find the attractors, then we must remove another dependency that was unmasked by the last equation: 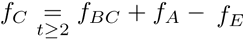, which implies three useful constraints.

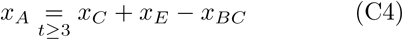

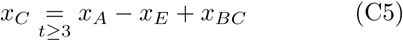

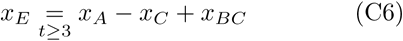

Each constraint reduces the size of the variable space: for example, the first eliminates all variables containing index A with no C, E, or BC. We did not solve for *x*_*BC*_ because doing so would not eliminate any variables, because the *x*_*C*_ term does not attach any new indices.

After applying the new constraints we obtain:

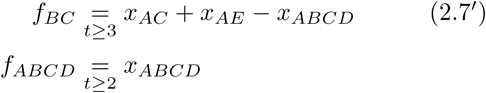

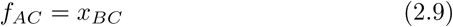

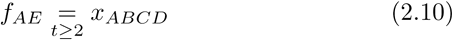

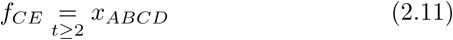

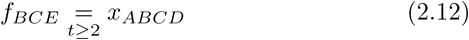

This produces new dependencies: 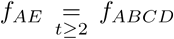 and 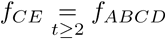, implying that

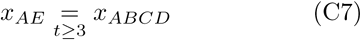

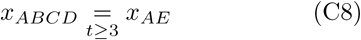

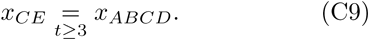

With these constraints our example ends with the following closed system of dynamical equations:

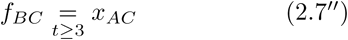

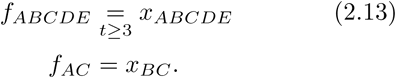

The attractor is always reached at or before time step 3. The constraint equations (C1-C9) map our variables of interest (*x*_*A*_, *x*_*C*_, and *x*_*E*_) to linear combinations of variables in the final system: for example, *x*_*A*_ is mapped by constraint C4 to *x*_*AC*_ + *x*_*AE*_ - *x*_*ABC*_, then mapped using constraints C1, C7, and C8 to *x*_*AC*_, whose dynamics are given by the final time-evolution equations. The eigenvalues of this final system are (-1, 1, 1), implying a steady state along with a period-2 cycle.

**Figure 4.**
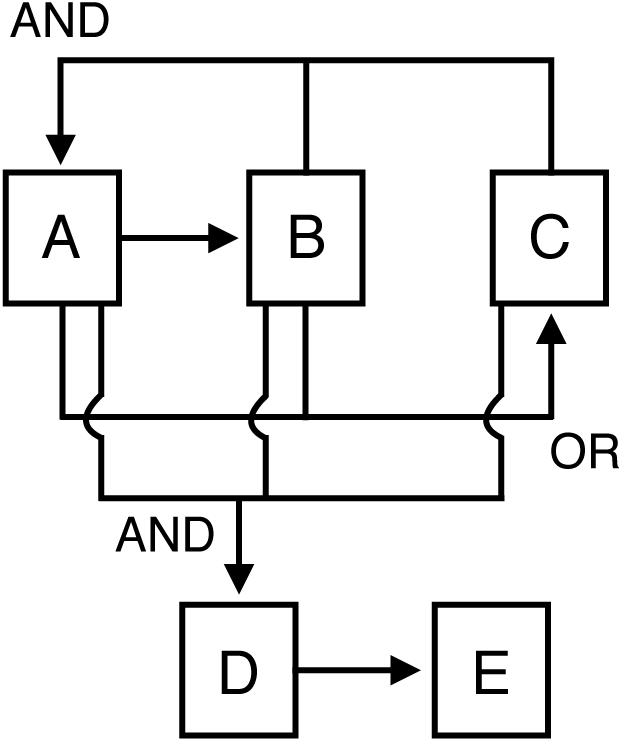
The network used in Example 2.

When using equation-reduction method 2, the order in which we calculate the time-evolution of new variables and remove dependencies greatly influences the complexity of the calculation. One general rule we have found is that constraints on the lowest-index variables are most helpful for deterministic networks, because they factorize the largest part of the variable space. Following this rule of thumb, our implementation solves the time-evolution equations for only those variables having the fewest indices between each search for new dependencies. Additionally, our code solves for low-index factors of new variables even if they were not directly involved in earlier equations in hopes that they will produce helpful constraints. In most tests, this prioritization method speeds up the calculation.

Equation-reduction method 2 is susceptible to numerical problems that fall into two categories: 1) polynomial coefficients can become very large, and 2) gradual erosion of numerical precision can blur the distinction between zero and nonzero terms. In addition our code monitors both sources of error; in the latter case by tracking the floating-point error in each step of the calculation. When either type of calculation error exceeds defined bounds, our method exits with an error message.

The reason polynomial coefficients can become large is that multiplication by constraints can produce several identical terms that add together. For example, the constraint *x*_1_ = *x*_2_ + *x*_3_ takes *x*_123_ → 2*x*_123_. Typically the variable being multiplied would be found to be zero later in the calculation (so having a coefficient of 2 is not an error in the math), but if many such constraints are applied and the coefficient grows exponentially then the numerics will eventually fail. We have included a filter that can usually identify these outlier coefficients and remove them using the rule that all individual (not population-averaged) product variables are either 0 or 1 at all time steps. For example, if *ƒ*_4_ = 2*x*_123_ then we can reason that if *ƒ*_4_ is either 0 or 1, then *x*_123_ cannot be 1 and therefore we have a new constraint *x*_123_ = 0 which simplifies *ƒ*_4_ → 0. In general, we identify these removable terms using the heuristic that if a large coefficient multiplying a combination of integer-weighted variables is greater than the sum of all other coefficients plus one, then that combination of variables (which is an integer for a homogeneous population) must be zero.

Our equation-reduction method 2 was tested by running 10^4^ comparisons between Monte Carlo and our dynamical equations using a random amount of equation-reduction (which determines the earliest time point at which the simulation is valid) for each test. We then compared the predictions of our exact simulations to the population dynamics as predicted by Monte Carlo. In each case the product-basis simulation exactly reproduced the average of the Monte Carlo simulations. These networks were small, but still provided a stringent test of the numerics, as they could theoretically involve linear systems of equations up to size (2^10^)^2^. During this test, our program also generated additional networks whose numerical error overstepped a threshold and were thus omitted from the Monte Carlo comparison; overall 18.5% of tests generated were discarded.

## Appendix 3: algorithm and code

Pseudocode is given in Algorithm 1. The full code is available at: https://github.com/CostelloLab/ProductBasis.

### Algorithm 1 build closed system of equations *F*

~~~
1: Initialize set of unsolved variables with variables of interest: *X* ← {*x*_1_, *x*_2_, …, *x*_*n*_}
2: Initialize set of variable update rules: 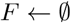
3: Initialize set of constraints: 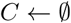
4: Initialize simulation start time: *t*_*sim*_ ← 1
5: Initialize set of equation start times of *F*: 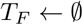
6: Initialize set of constraint start times: 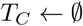

7: **while** *X* is not empty **do**

8:    % Reduce equations if necessary
9:    **if** *size*(*F*) > *equation_reduction_threshold* **then**
10:       *F*_□_ ← square-matrix component of *F* (i.e. only terms in solved variables for each *ƒ*_*i*_)
11:       λ ← set of eigenvalues of 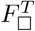
12:       **for** each λ_i_ ∈ λ **do**
13:          *G* ← *F* – λ_i_[*I* 0]
14:          *R* ← upper-triangular matrix from *QR*(*G*)
15:          *V*_*j*_ ← set of dependent variables *j* found from |*R*_*jj*_| < *ϵ*
16:          *D* ← set of linear-dependencies found by 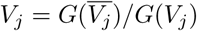
17:          **if** 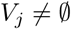 **then**
18:                  *t*_*sim*_ ← min(*t*_*sim*_, min(*T*_*F*_(*x*_*k*_ ∈ *D*)) + max(1, ceiling(log *ϵ*_*decay*_/log |λ_*i*_|)))
19:                  **break**
20:                **end if**
21:             **end for**
22:             *D* ← sort {*d*_*i*_ for which *n*_*terms*_(*d*_*i*_) ≤ 2min *n*_*terms*_(*d*)} by number of terms
23:             **for** each *d*_*i*_ ∈ *D* **do**
24:                 NewConstraints(*d*_*i*_, *t*_*sim*_)
25:               **end for**
26:               Sort *C* by number of terms
27:             **end if**

28:             % Add new equations
29:             *X* ← *X* ⋃ lowest-index factors of *X*
30:             Sort *X* by number of terms
31:             **for** each *x*_*i*_ ∈ *X* **do**
32:                 *ƒ*_*i*_ ← 1
33:                 **for** each Boolean factor *x*_*b*_ of *x*_*i*_ **do**
34:                    **if** *ƒ* is a discrete update rule **then**
35:                        *ƒ*_*i*_ ← *ƒ*_*i*_ · *ƒ*_*b*_
36:                     **else**
37:                        *ƒ*_*i*_ ← *ƒ*_*i*_ + *ƒ*_*b*_ · ∏_*x*_*j*_ ≠*x*_*i*_ ∈*X*_ *x*_*j*_
38:                     **end if**
39:                   **end for**
40:                   *F* ← *F* ⋃ Constrain(*ƒ*_*i*_, 1)
41:                 **end for**

42: **end while**
43: **function** NewConstraints(*d*, *t*_*c*_)
44:    **if** *d* ← RemoveOversizeCoefficients(*d*/minCoef(*d*)) adds no new constraint **then**
45:       **for** each *x*_*j*_ ∈ *d* that does not factor another *x*_*k*≠*j*_ ∈ *d* **do**
46:          *RHS*_*j*_ ← solve (*d* = 0) for *x*_*j*_
47:          **if** {*x*_*j*_} ≠ Constrain({*RHS*_*j*_}, *t*_*c*_)[1] **then**
48:              **for** each prior constraint (*x*_*p*_ = *RHS*_*p*_) **do**
49:                  **if** *x*_*p*_ is factorized by *x*_*j*_ **and** *x*_*p*_ · *RHS*_*j*_ = *RHS*_*p*_ **then**
50:                      *C* ← *C* \ {*c*_*p*_}
51:                    **end if**
52:                 **end for**
53:                 *C* ← RemoveOversizeCoefficients(*C* ⋃ {*x*_*j*_ = *RHS*_*j*_})
54:                 **for** each *x*_*k*_ = multiples of *x*_*j*_ **do**
55:                     *X*_*new*_ ← new terms in Constrain(*x*_*k*_)
56:                     *X* ← *X* ⋃ *X*_*new*_
57:                     *A* ← {*a*_*l*_ = coefficients of *x*_*k*_ in *F*}
58:                     *F* ← *F* + *A* · (*RHS*_*k*_ - *x*_*k*_)
59:                     (*t*_*ƒ*_*l*__ for all *l* such that *a*_*l*_ ≠ 0) ← *t*_*c*_
60:              **end for**
61:           **end if**
62:        **end for**
63:     **end if**
64: **end function**
65: **function** Constrain(*poly*, *t*_*c*_)
66:     **while** ∃ unconstrained terms in *poly* **do**
67:       **for** each constraint *x*_*i*_ = *RHS*_*i*_ **do**
68:          % Multiply all factored variables by the constraint
69:          **for** each unconstrained variable *x*_*j*_ with coefficient *a*_*j*_ in *poly* **do**
70:             **if** *x*_*j*_ is factorized by *x*_*i*_ **then**
71:                *poly*_*new*_ ← *x*_*j*_ · *RHS*_*i*_
72:                **if** *x*_*j*_ ⊄ *poly*_*new*_ **then**
73:                    *poly* ← (*poly* ← *a*_*j*_ · *x*_*j*_) ⋃ *poly*_*new*_
74:                    *t*_*ƒ*_*i*__ ← *t*_*c*_
75:                 **end if**
76:              **end if**
77:            **end for**
78:            *poly* ← RemoveOversizeCoefficients(*poly*)
79:         **end for**
80:      **end while**
81:      **return** *Polys*
82: **end function**

83: % Check for overlarge coefficients
84: **function** RemoveOversizeCoefficients(*poly*)
85:    **while** ∃ unchecked coefficients in *poly*_*i*_ **do**
86:      **for** each coefficient *a*_*i*_ in *poly*_*i*_ **do**
87:          *n*_*j*_ = round(*a*_*j*_/*a*_*i*_)
88:          **if** |*a*_*i*_| > ∑_*j*_ |*a*_*j*_ - *n*_*j*_*a*_*i*_| **then**
89:              NewConstraints(*n*_*j*_, *t*_*sim*_)
90:              *a*_*j*_ ← *a*_*j*_ - *n*_*j*_*a*_*i*_ for all *j*
91:              **break**
92:           **end if**
93:        **end for**
94:      **end while**
95:      **return** *Polys*
96: **end function**
~~~

## Acknowledgments

We would like to thank Sharon Lutz for giving useful mathematical advice, and Federico Stefanini and Julio Saez-Rodriguez for helpful comments on the manuscript. This work is supported by the Boettcher Foundation (J.C.C.) and NIH grant 2T15LM009451 (B.C.R.).

